# Viral protease-mediated polyprotein processing in human astroviruses

**DOI:** 10.1101/2025.11.24.690148

**Authors:** David Noyvert, Leandro X. Neves, Ksenia Fominykh, Jacqueline Hankinson, Gemma Lindsey, Aleksei Lulla, Edward Emmott, Valeria Lulla

## Abstract

Positive-sense RNA viruses often encode large polyproteins that are proteolytically processed by viral and host proteases into functional replication proteins. Astroviruses infect intestinal and neuronal cells across diverse human and animal hosts and also rely on polyprotein cleavage for replication. In this study, we mapped the cleavage sites of the nonstructural polyproteins of classical human astrovirus 1 (HAstV1) and neurotropic astrovirus strain MLB2 using complementary N-terminomics of infected cells and analyses of untagged overexpressed polyproteins. Notably, we identified two adjacent cleavage sites at the N-terminus of HAstV1 and MLB2 proteases, as well as a similar dual cleavage site at the C-terminus of the MLB2 protease. We also demonstrated processing of the hypervariable region and VPg in both astrovirus strains. This allowed us to define the boundaries of individual protein products and identify conserved and divergent processing features between classical and non-classical astroviruses. Additionally, we characterized several polyprotein precursors and evaluated the replication properties of cleavage-deficient mutant replicons, revealing the critical role of polyprotein processing for functional replication complex formation. Understanding the dynamics of polyprotein processing is essential for interpreting the stages of viral infection and identifying new drug targets and antiviral strategies.

## Introduction

Astroviruses are non-enveloped positive-strand RNA (+ssRNA) viruses that infect mammals (*Mamastrovirus*) and birds (*Avastrovirus*), causing a variety of mild to severe pathologies. They are common causes of gastrointestinal disease in the human population, with most infections spread through the fecal-oral route. Seroprevalence studies have shown that up to 90% of adults have been exposed to astrovirus infection^1,2^. Classical astroviruses (HAstV1-8) are associated with mild, self-limiting gastrointestinal symptoms such as diarrhea. However, newly discovered non-classical astroviruses from MLB and VA/HMO genogroups present extra-intestinal pathogenicity, with several documented cases of meningitis and encephalitis in the immunocompromised and elderly^3,4^. There are currently no treatments or vaccines against astroviruses, highlighting the importance of understanding their basic biology to mitigate the impact of future outbreaks.

Astroviruses have a short genome of 6.8-8.7 kb arranged in three or four open reading frames (ORFs), flanked by 5′ and 3′ untranslated regions. Following entry into the host cell, mediated by binding to Fc and DPP4 receptors and subsequent endocytosis^5^, the 5′ VPg-linked viral genome is uncoated in the cytoplasm and translated into full-length nonstructural protein 1a (nsP1a). This polyprotein is subsequently processed into individual proteins, including the N-terminal domain (NTD), transmembrane (TM) helices, viral protease (Pro), viral protein genome-linked (VPg), and hypervariable region (HVR)^6,7^. A programmed ribosomal frameshift between nsP1a and nsP1ab enables translation of viral RNA-dependent RNA polymerase (RdRp) from nsP1ab polyprotein^8^. Additionally, two ORFs are expressed from subgenomic RNA: ORF2 encodes the capsid polyprotein, and ORFX (present in most astroviruses) encodes XP, a viroporin^9^.

The NTD contains di-arginine motifs that target the polyprotein to the perinuclear endoplasmic reticulum (ER) membrane, facilitating the assembly of the viral replication complex. Both cellular and viral proteases process the polyprotein to release proteins essential for the replication of the viral genome^10^. The astrovirus protease is a serine protease with a previously determined crystal structure^11^. It has an Asp-His-Ser catalytic triad that recognizes glutamine residues during proteolytic processing. Mutation of the serine residue of the catalytic triad abolishes catalytic activity^12^. Although the mechanistic role of VPg in the astrovirus life cycle has not been experimentally confirmed, its function can be inferred from the role of VPg in caliciviruses, where it is thought to attract translation initiation factors such as eIF4 and eIF3^13–15^. Previous studies have shown that VPg is essential for viral infectivity since treatment of astrovirus RNA with proteinase K abolished the recovery of infectious virus^16^. The C-terminal product of nsP1a processing contains HVR, and its role remains to be characterized. Previous studies of this nsP1a region have contained the VPg and suggested a role in viral RNA synthesis^17^. It is possible that the function of the fully cleaved HVR may be different. Genetic analysis of the C-terminus of nsP1a from clinical samples revealed an association between this region and the pathogenicity and severity of gastroenteritis^18^.

The exact boundaries of the processing and the function of all the products are still unknown. The number of cleaved proteins also likely exceeds the current terminology, suggesting that four proteins are formed during the cleavage of nsP1a (nsP1a/1-nsP1a/4). We employ domain-based protein nomenclature to mitigate this discrepancy in the current study. Polyprotein processing is a key step in the life cycle of +ssRNA viruses, liberating proteins for membrane rearrangements and genome replication. Astrovirus polyprotein processing is described in the protein database UniProt, which predicts processing patterns of nsP1a of HAstV1^19^. However, these sites have not been experimentally confirmed in virus-infected cells. *In vitro* translation studies have demonstrated that HAstV1 nsP1a can self-cleave to produce 64 and 38 kDa products from the 101 kDa nsP1a polyprotein^12^. Analysis of cleavage products from HAstV1-infected Caco-2 cells revealed proteins of 75, 34, 20, and 8 kDa^20^. Other studies have observed final cleavage products of 57, 27, and 20 kDa corresponding to the putative RdRp, protease, and N-terminal domain, respectively^21,22^. Additionally, in a recently published work, Mehri *et al.* used tagged overexpressed HAstV1 polyproteins to identify cleavage sites in nsP1a/b that partially overlapped with those predicted by Uniprot and other sources. However, these cleavages were not validated in astrovirus-infected cells^23^.

In this study, we performed proteomic analyses of cells infected with two genetically diverse astrovirus strains, HAstV1 and MLB2, and complemented by analysis of untagged overexpressed polyproteins in mammalian cell lines. This approach enabled us to map the polyprotein processing of classical HAstV1 and non-classical MLB2 nsP1a by viral protease. These cleavage maps expand the repertoire of previously reported cleavage sites and provide a platform for comparative analysis of the nsP1a cleavage pattern between two distinct astrovirus strains. To assess the functional significance of these cleavage events, we employed HAstV1-and MLB2-based subgenomic replicon systems, demonstrating that precise polyprotein processing is essential for efficient astrovirus RNA replication. Collectively, our findings define the map of HAstV1 and MLB2 astrovirus protease-mediated cleavage products and establish a foundation for the development of protease-targeting therapeutics.

## Results

### Detection of ORF1a polyprotein cleavage products in HAstV1– and MLB2-infected cells

The ORF1a-encoded polyprotein is presumably cleaved into individual proteins, referred to as NTD, TM, Pro, linker (L), VPg, and HVR (Fig. 1A). However, there has been minimal characterization of individual products, their boundaries, and intermediates in astrovirus-infected cells. To comprehensively address this question, we generated a panel of antibodies to detect astrovirus proteins and their precursors: HAstV1 VPg, HAstV1 HVR, and MLB2 VPg, as well as previously described HAstV1 Pro and MLB2 Pro^10^. The full HAstV1 VPg and HVR, as well as the N-terminal part of MLB2 VPg possessing a C-terminal 8×His-tag were used for bacterial expression and affinity purification, resulting in homogeneous proteins of the expected sizes (Fig. 1B). Each purified recombinant protein was used for the production and affinity-based purification of antibodies^10,24^.

**Figure 1.**
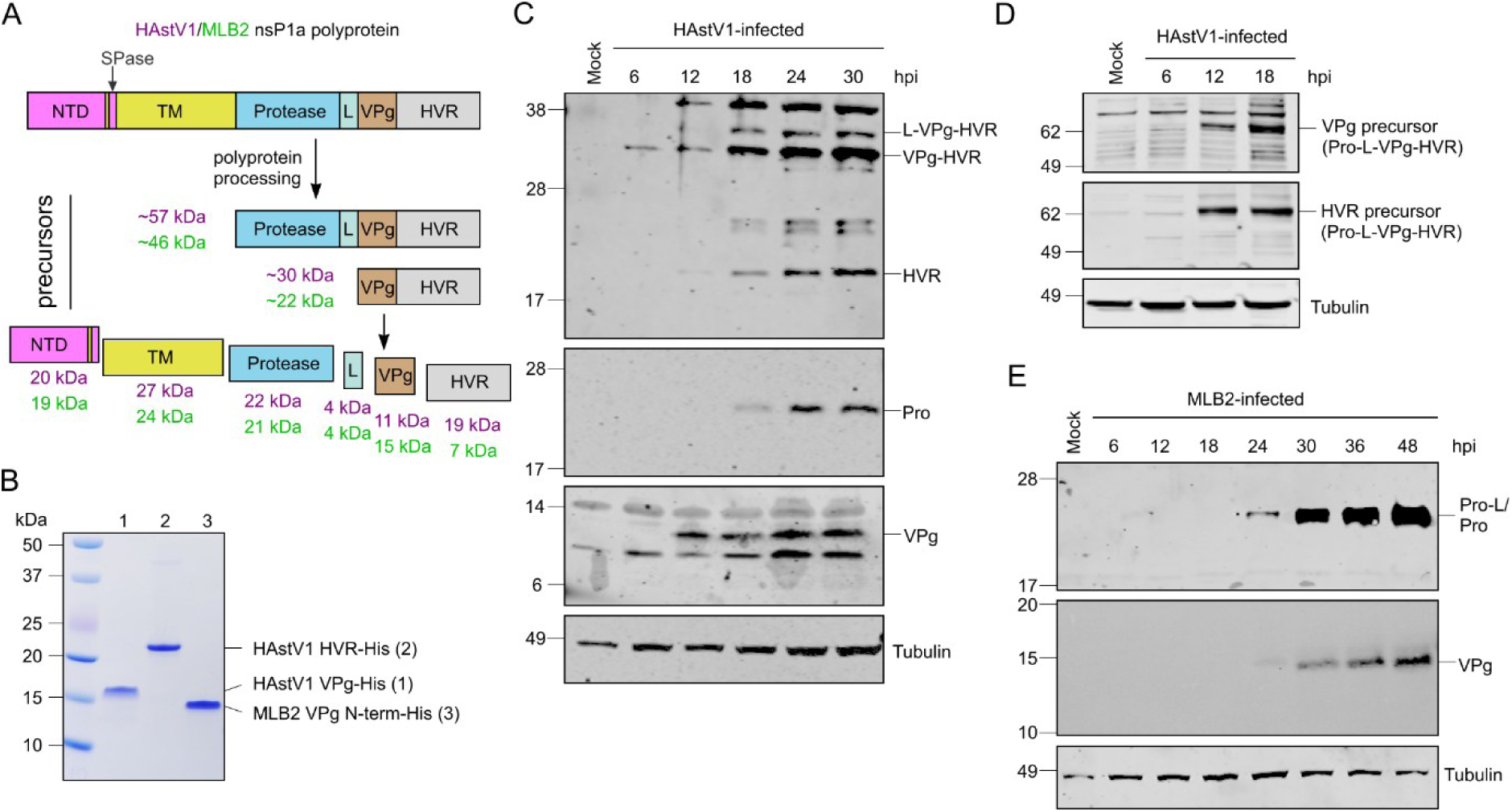
Detection of HAstV1 and MLB2 astrovirus proteins during infection. (**A**) Schematic representation of HAstV1 and MLB2 nonstructural polyprotein nsP1a, predicted cleavage products, and their precursors. NTD, N-terminal domain; TM, transmembrane region; L, 4 kDa linker; VPg, virus protein genome-linked; HVR, hypervariable region. (**B**) Coomassie-stained SDS-PAGE profile of the purified HAstV1 VPg and HVR and MLB2 VPg (N-terminal part) from *E. coli*. HAstV1 and MLB2 Pro were described previously^10^. (**C**) Caco-2 cells were infected with HAstV1 at an MOI 1, harvested at indicated time points, and analyzed by western blotting using custom anti-HAstV1 HVR, Pro, and VPg antibodies. (**D**) The same samples (C) were analyzed for the presence of precursor proteins detected using anti-HVR and anti-VPg antibodies but at different exposure times. (**E**) Huh7.5.1 cells were infected with MLB2 at an MOI 1, harvested at indicated time points, and analyzed by western blotting using custom anti-MLB2 antibodies against Pro and VPg.

The detection of cleavage products was performed to understand the dynamics of nsP1a processing and identify viral proteins produced during infection in Caco-2 (HAstV1) and Huh7.5.1 (MLB2) cell lines. Following HAstV1 infection, a ∼22 kDa viral protease and a ∼20 kDa HVR were detected at 18 hours postinfection (hpi), corresponding to the predicted protein sizes. VPg was detected at 12 hpi, with the predicted protein size of ∼11 kDa, and levels remained constant throughout the infection (Fig. 1C). The detection of ∼60 kDa precursor protein was possible from 12 hpi with VPg– and HVR-detecting antibodies (Fig. 1D). This precursor likely corresponds to Pro-L-VPg-HVR (Fig. 1A). Despite good detection, antibodies against Pro did not detect this ∼60 kDa precursor, suggesting potential differences in the immunodetection of individual and unprocessed protease forms. The cleavage of the N-terminal domain was previously investigated using NTD-HA-tagged HAstV1 and MLB2 astroviruses, demonstrating the detection of processed NTD-HA at 18 hpi (HAstV1) and 24 hpi (MLB2)^10^. Consistent with its slower replication rate^24^, the processing of MLB2 polyprotein resulted in the detection of the ∼21 kDa protease and ∼15 kDa VPg products from 24 hpi (Fig. 1E).

Taken together, we detected the following processed HAstV1 proteins: NTD, Pro, VPg, and HVR, confirming previously suggested processing patterns. For MLB2, the cleavages of NTD, Pro, and VPg were also confirmed, suggesting the cleavage of the C-terminal HVR. Their exact boundaries, roles, and dynamics of the cleavage remain to be characterized.

Based on these analyses, we selected 20 hpi for HAstV1– and 40 hpi for MLB2-infected cells to prepare samples for N-terminomic analyses, aiming to detect the exact boundaries of proteolytically processed viral proteins in virus-infected cells.

### Identification of neo-N-termini generated through protease activity using proteomic analysis of HAstV1– and MLB2-infected cells

N-terminomic analysis of HAstV1– and MLB2-infected cells was performed to investigate proteolysis during astrovirus infection. This approach relies on the detection of novel N-termini during virus infection via protease cleavage of astrovirus polyproteins (Fig. 2A). Due to the length and properties of generated peptides, the mode of sample preparation, and detection limits, some cleavage sites may still remain undetected. We, therefore, relied on the data combined from both viruses to create a map of cleavage sites for the follow-up analyses.

**Figure 2.**
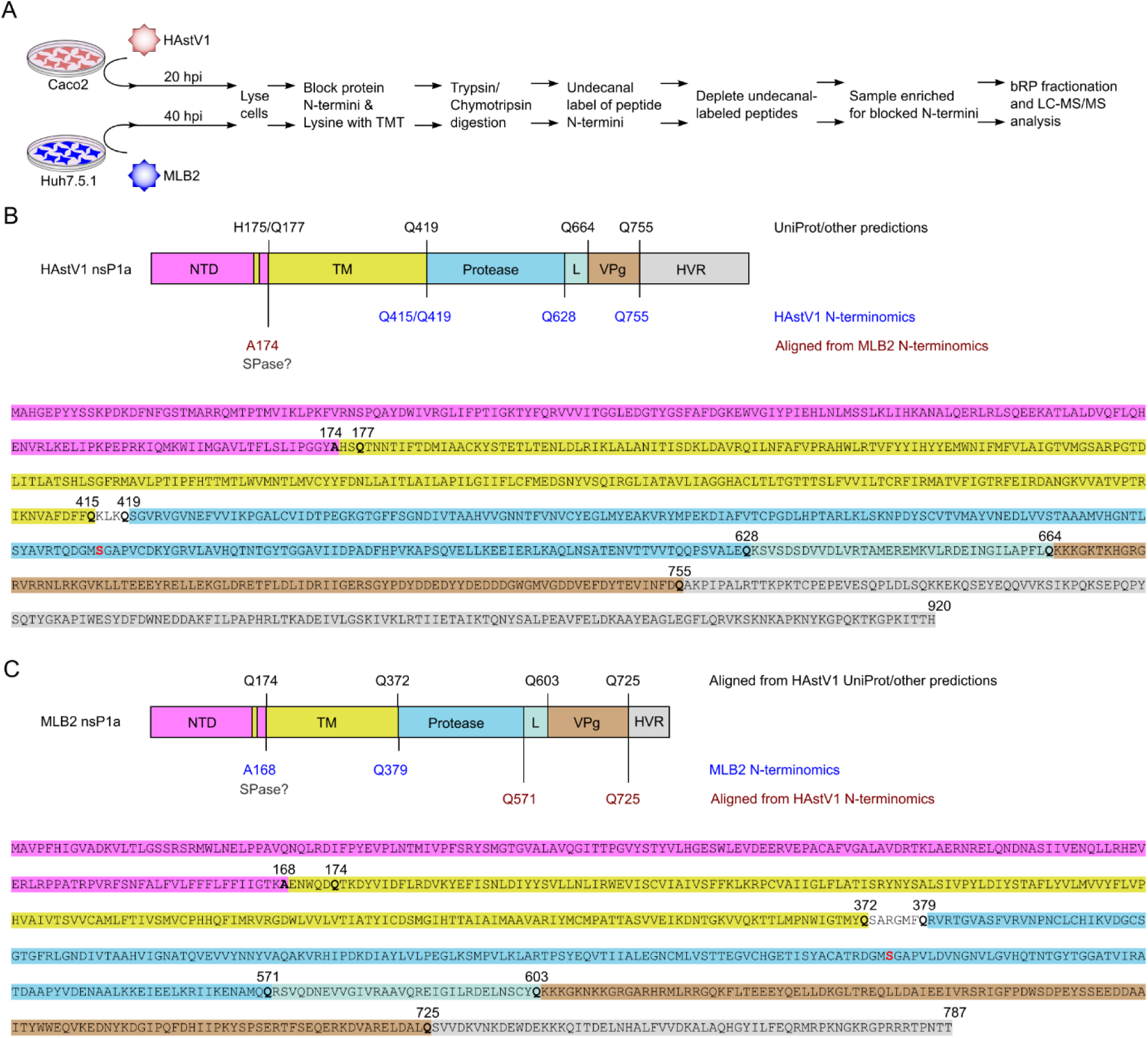
Proteolysis of viral proteins during astrovirus infection. (**A**) Experimental design of N-terminomic analysis. (**B-C**) Schematic of the HAstV1 (B) and MLB2 (C) astrovirus nsP1a with UniProt or otherwise predicted (black), identified through N-terminomics analysis (blue) or predicted based on homology between HAstV1 and MLB2 (maroon) cleavage sites. The catalytic serine protease residue (S) is marked in red (B-C).

First, we compared the cleavage sites previously predicted by UniProt^19^ and obtained in proteomic analyses for two astroviruses, HAstV1 and MLB2. For HAstV1, the predicted processing sites by UniProt were H175, Q419, Q664, and Q755 (UniProt entry Q67726). We added Q177 as the potential viral protease cleavage-mediated site. N-terminomics analysis revealed novel sites at Q628 and Q415, and confirmed the predicted Q419 and Q755 sites (Fig. 2B). Based on homology with HAstV1 predictions, the potential processing sites in MLB2 were Q174, Q372, Q603, and Q725. The experimentally detected sites were mapped to A168 and Q379 (Fig. 2C). The full list of nonstructural polyprotein-derived peptides detected in proteomic analyses is provided in Table S1, excluding peptides with trypsin– and chymotrypsin-generated N-termini. We compared cleavage patterns revealed by proteomic analysis to predict missing boundaries for the HAstV1 and MLB2 polyprotein processing (Fig. 1C-E). Both predicted and newly identified serine protease cleavage sites were selected for further validation. The cleavage after A168 in MLB2 is likely a cellular signal peptidase (SPase) cleavage product^10,25^, corresponding to the HAstV1 analogous position at A174 (Fig. 2B-C). The Q526 cleavage (Table S1) was not investigated, as it does not correspond to the observed protease-containing cleavage product (21 kDa, Fig. 1E).

### Mutational analysis of viral protease-mediated proteolytic processing of HAstV1 astrovirus nonstructural polyprotein

To investigate predicted and detected cleavage sites (Fig. 2), full-length HAstV1 nsP1a sequence was cloned into a mammalian expression vector (Fig. 3A) and overexpressed in HEK293T cell line. This cell line supports replication of astroviruses^10,24^ and, therefore, was considered suitable for studying polyprotein processing. Mutations at the predicted and detected cleavage sites were introduced into the nsP1a expression vector to investigate the effect on polyprotein cleavage patterns. The glutamine residue was mutated to glycine to prevent cleavage at the P1 residue by the viral protease.

**Figure 3.**
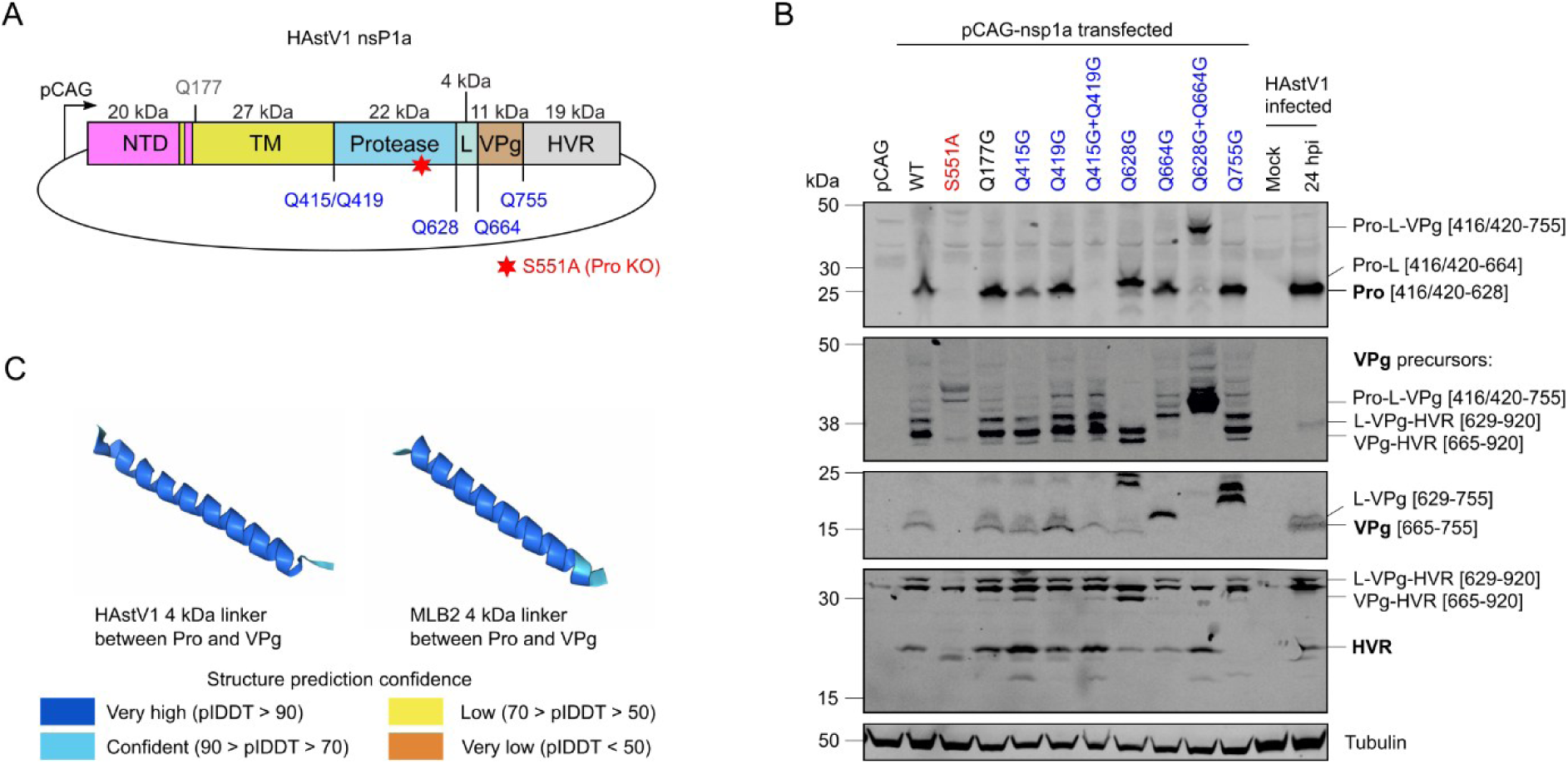
Investigating processing of the overexpressed HAstV1 nsP1a. (**A**) Schematic representation of pCAG mammalian expression vector used to overexpress nsP1a of HAstV1. (**B**) HEK293T cells were transfected with pCAG plasmids expressing wt and mutant nsP1a of HAstV1 and analyzed by western blotting using HAstV1 anti-Pro, anti-VPg, anti-HVR, and human anti-tubulin antibodies. Lysate obtained from HAstV1-infected Caco-2 cells was used as a control. (**C**) 3D model for HAstV1 and MLB2 4 kDa linker between protease and VPg predicted with AlphaFold 3.

As expected, the products of wild type (wt) nsP1a expression and subsequent cleavage corresponded to those derived from infected cells (Fig. 3B). The protease knockout mutant S551A was used as a cleavage-deficient control (Fig. 3A-B). The cleavage of NTD was not blocked by the predicted Q177 mutation, indicating that this cleavage is unlikely to be performed by the viral protease, which cleaves at glutamine residues, and confirming previously suggested cleavage by cellular protease(s)^24^.

The cleavage at Q415 and/or Q419 is supposed to release the N-terminus of HAstV1 protease. Individually, Q415G and Q419G did not abolish polyprotein processing. However, combining these mutations prevented proteolytic processing of the viral protease (Fig. 3B), suggesting redundancy of cleavage sites for the release of the N-terminus of the HAstV1 protease.

The processing at Q628 should result in a protease-containing product of ∼22 kDa, observed in HAstV1-infected and pCAG-nsP1a transfected cells (Fig. 3B). The mutation Q628G resulted in ∼4 kDa larger protease product, suggesting another closely located interdomain cleavage site, likely representing the linker between K629 and Q664 (predicted molecular weight of 4 kDa). The mutation Q664G restored the correct protease processing pattern but increased the size of the VPg, suggesting the N-terminal boundary of this protein. Mutation of both Q628 and Q664 resulted in the abolishment of protease and VPg processing, creating longer products that were strongly detected using VPg– and Pro-specific antibodies. The role of ∼4 kDa cleaved peptide between Pro and VPg remains to be elucidated. Interestingly, its structure is predicted as a helix with high confidence in both HAstV1 and MLB2 (Fig. 3C). The mutation of Q664 confirms the cleavage site for the N-terminus of the VPg. The detected processing product is ∼4 kDa larger, corresponding to the K629-Q755 VPg-containing product (Fig. 3B).

The final HAstV1 nsP1a cleavage site predicted by UniProt and detected by N-terminomics (Fig. 2A-B) at Q755 should separate VPg and HVR. Indeed, the Q755G mutation increased the size of both proteins, suggesting the correct cleavage site. The additional ∼20 kDa bands appearing on the VPg-probed blot suggest potential alternative cleavages within HVR when Q755 is blocked. The appearance of smaller HVR-specific bands (lanes 5, 7, 10, 11) suggests alternative cleavage patterns when corresponding cleavage sites are blocked (Fig. 3B). With the abolishment of the protease cleavage at Q415 and Q419, the VPg and HVR can still be processed (Fig. 3B). This indicates that the protease precursors may still retain proteolytic activity. Mehri *et al.* recently identified cleavage sites within HAstV1 nonstructural polyprotein using overexpressed dual-tagged constructs. Their findings also confirmed cleavage at sites Q415, Q628, Q664, and Q755 in HAstV1 nsP1a, although they did not detect cleavage at Q419^23^.

### Mutational analysis of viral protease-mediated proteolytic processing of MLB2 astrovirus nonstructural polyprotein

The processing of MLB2 was investigated using overexpressed MLB2 nsP1a polyprotein. The protease knockout mutant S510A was used as a cleavage-deficient control (Fig. 4A-B). Similar to the HAstV1 Q177 mutant, the cleavage of the NTD was not blocked by Q174 in MLB2 polyprotein, further suggesting cleavage by cellular protease(s) at this site^24^. However, there was a decrease in efficiency of Pro and VPg processing, suggesting an impact on cleavage efficiency (Fig. 4B), possibly due to interference with NTD processing at A168 (Fig. 2C) and/or membrane association.

**Figure 4.**
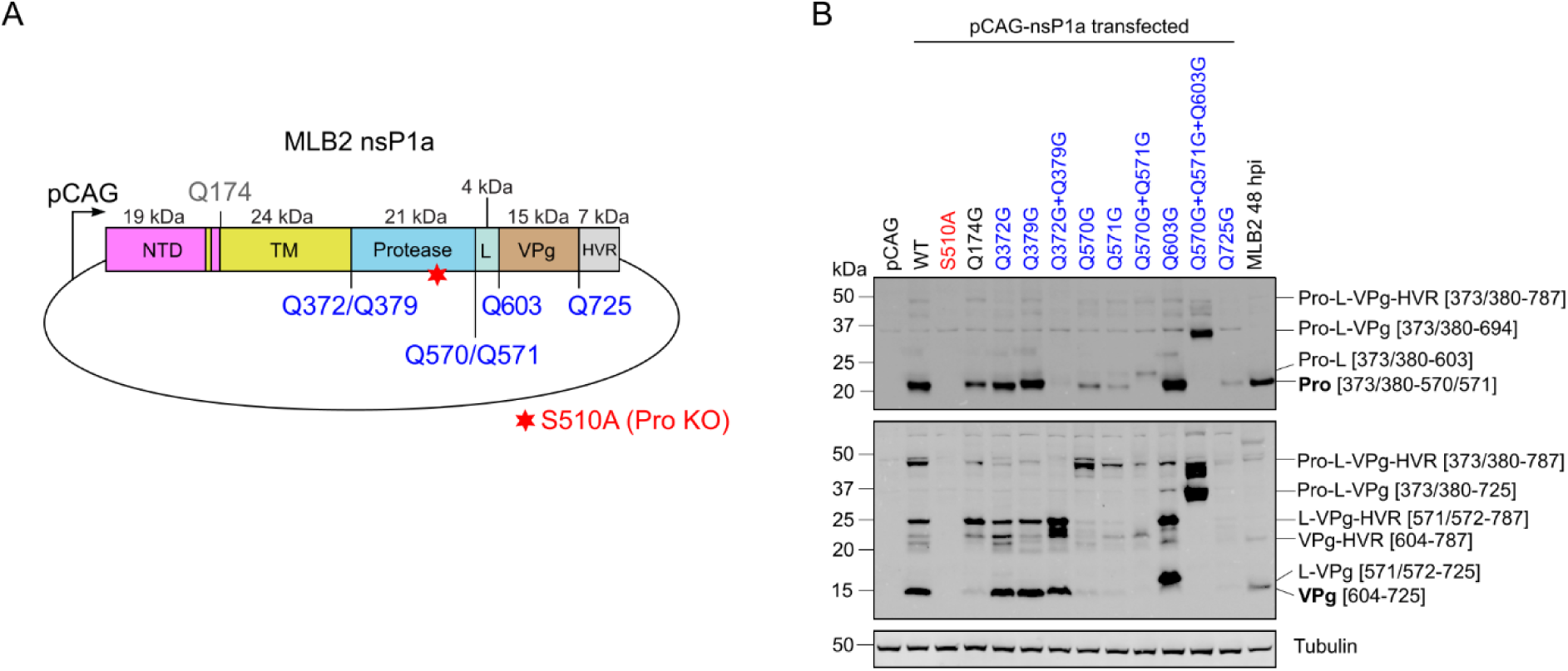
Investigating processing of the overexpressed MLB2 nsP1a. (**A**) Schematic representation of pCAG mammalian expression vector used to overexpress nsP1a of MLB2. (**B**) HEK293T cells were transfected with pCAG plasmids expressing wt and mutant nsP1a of MLB2 and analyzed by western blotting using custom MLB2 and anti-tubulin antibodies. Lysates obtained from MLB2-infected Huh7.5.1 cells were used as a control.

Based on the HAstV1 N-terminomics data, the N-terminus of MLB2 protease can be cleaved at Q372 and Q379. Mutating these residues individually did not affect the release of the Pro N-terminus (Fig. 4B). Similarly to HAstV1, mutation of the two residues together blocked the release of fully processed Pro. In contrast to HAstV1, there was also redundancy in cleavage sites at the C-terminus of the MLB2 protease. Mutating the Q571 residue alone reduced the Pro intensity but did not fully disrupt the processing, suggesting that the protease can compensate for this cleavage using position Q570. The mutation of Q570 and Q571 together resulted in a larger protease band, likely up to the residue Q603 (Fig. 4B). The mutation of all three sites together (Q570, Q571 and Q603) prevented corresponding cleavages, resulting in the detection of ∼37 kDa protein (Fig. 4B), likely Pro-L-VPg precursor that is not characteristic of wt nsP1a processing.

Using MLB2 VPg-detecting antibodies, we investigated the C-terminal polyprotein cleavages. Similarly to HAstV1, mutating the Q372 and Q379 dual residues at the N-terminus of protease did not affect the release of fully cleaved VPg. This suggests that the protease precursors have proteolytic activity. Mutating the C-terminal protease cleavage site has a greater effect on VPg processing efficiency. Q603G mutation results in detection of a larger VPg product due to cleavage at Q570/Q571. Mutating Q570, Q571 and Q603 together abolishes VPg processing and results in the accumulation of ∼37 kDa Pro-VPg precursor that can be detected by both anti-Pro and anti-VPg antibodies (Fig. 4B). These experiments further confirm cleavage at Q571, Q603, and minor cleavage at Q570. Mutation of C′ of MLB2 VPg at Q725 prevented VPg processing, suggesting this site is the boundary between VPg-HVR.

These experiments confirmed the following processing sites in MLB2 nsP1a: Q372, Q379, Q571, Q603, Q725, and minor cleavage at Q570.

Taken together, our data demonstrate differences in the processing of classical (HAstV1) and non-classical (MLB2) nsP1a. According to the presented data (Figs 3 and 4), processing of HAstV1 and MLB2 nsP1a should yield six cleavage products. However, they differ in size and redundant cleavage patterns.

### Impact of cleavage site mutations on viral replication using HAstV1 and MLB2 replicon systems

Cleavage of the astrovirus nonstructural protein is a crucial step in virus replication, enabling the correct localization and formation of the replication complex^10^. To investigate the importance of detected cleavage sites in the context of viral replication, the set of mutants (Figs 3 and 4) was introduced to HAstV1 and MLB2 replicons. In the previously described HAstV1– and MLB2-based replicon systems^9,24^, RNA replication is evaluated via a luminescent reporter gene (Renilla luciferase) that replaces the structural proteins, allowing over two logs of dynamic range in replicon activity measurements (Fig. 5A). The controls for replication-deficient variants included RdRp knockout (GDD→GNN)^10,24^ and serine protease knockout (S→A). Inactivation of these main enzymatic units in both replicons resulted in >99% reduction of replicon activity when compared to the wild type. The replicon activity was measured in two cell lines to account for cell-specific effects (Fig. 5B-C).

**Figure 5.**
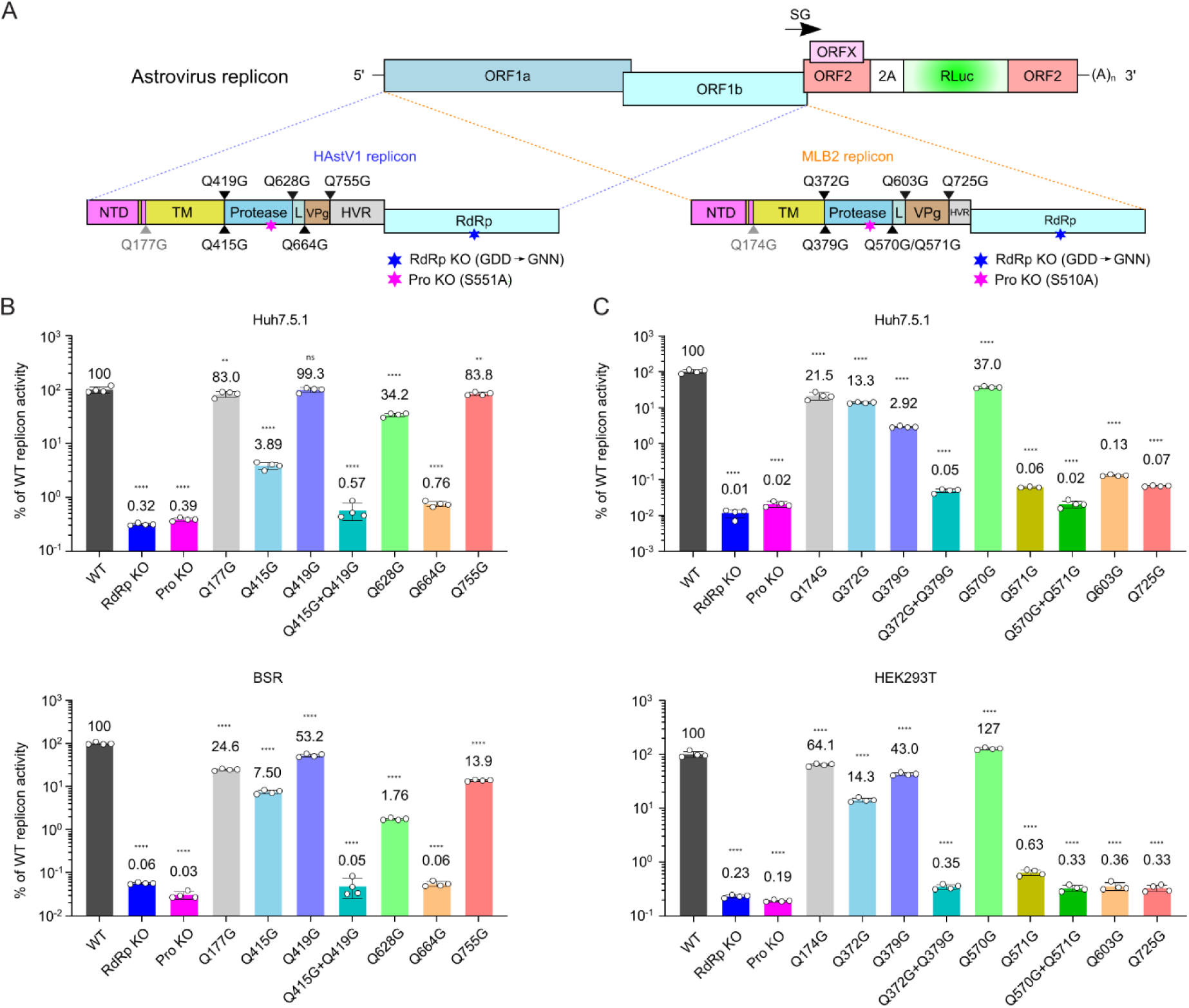
Polyprotein processing defines replication activity in HAstV1 and MLB2 replicon systems. (**A**) Schematic representation of HAstV1 and MLB2 replicons and introduced mutations. (**B**) Relative luciferase activities of HAstV1 replicons in Huh7.5.1 and BSR cells. (**C**) Relative luciferase activities of MLB2 replicons in Huh7.5.1 and HEK293T cells. The mean value is shown above each bar. Data are mean ± SD (*n* = 4, ≥3 independent experiments, graphs B-C). \*\**p* < 0.01, \*\*\**p* < 0.0001, ns, nonsignificant, using one-way ANOVA test against wt replicon.

HAstV1 replicons with Q419G mutation resulted in 1% (Huh7.5.1) and 47% (BSR) reduction of replicon activity (Fig. 5B), suggesting that these changes can be tolerated in the replicon context. This could be explained by redundant processing sites, as shown for Q415/Q419-mediated cleavage (Fig. 3B). Mutation of the Q415 processing site resulted in >90% decrease in replicon activity in both cell lines. Combining both Q415 and Q419 cleavage site mutants further reduced replicon activity to levels comparable to replication-deficient controls (>99%), highlighting possible redundancy of 415 and 419 cleavages (Fig. 5B). Mutating Q628, the C-terminal cleavage of protease from polyprotein, resulted in 64% (Huh7.5.1) and 98% (BSR) reduction of replicon activity. Mutating the Q664 residue at the N-terminus of the VPg resulted in >99% reduction of replicon activity, suggesting that the VPg may not be functional unless it is processed at the N-terminus. The tolerance of Q755-blocked cleavage (84% in Huh7.5.1 and 14% in BSR) is intriguing, considering the complete VPg-HVR cleavage block introduced by this mutation (Fig. 3B). This may imply partial functionality of VPg-HVR fused protein or its truncated versions.

The processing of the N-terminus of the MLB2 protease (Q372 and Q379) varied between cell lines, resulting in 57-97% reduction in replicon activity (Fig. 5C). Combined, these mutations abolish replication close to the replication-deficient control (Fig. 5C), confirming similar observations for HAstV1 replicon (Fig. 5B). Mutating Q570, which was shown to be a minor processing site, had a minor effect on replication. This is consistent with the overexpression data, which show no impact on protease or VPg processing. Mutating Q571 alone resulted in >99% reduction in replicon activity. Combining both Q571G and Q570G mutations further reduced replicon activity, suggesting their cumulative effect. Mutating Q603 and Q725 resulted in a significant decrease in replicon activity, comparable to the levels observed in replicase knockout controls, highlighting the importance of correct Pro and VPg processing for viral replication. In both HAstV1 and MLB2 replicons, the non-confirmed predicted cleavage of the NTD (Q177 in HAstV1 and Q174 in MLB2) resulted in 17-78% reduction of replicon activity (Fig. 5), suggesting viral protease cleavage-independent defects, likely due to the proximity of the TM region.

Overall, this data shows the importance of correctly processed sites for viral replication, highlighting some differences in processing requirements between classical and non-classical astroviruses. Between cell lines, we observe some differences in absolute numbers; however, the tendencies are well-reproducible.

### A consensus motif for HAstV1 and MLB2 protease cleavage sites

To determine a conserved cleavage motif for nsP1a cleavage sites, we analyzed representative sequences of closely related astrovirus species. The analysis revealed a conserved cleavage motif characterized by a glutamine residue (Q) at the P1 position and a hydrophobic residue (F, L, M, A) at the P3 position. This suggests a shared cleavage specificity among different regions of the astrovirus nonstructural polyprotein (Fig. 6).

**Figure 6.**
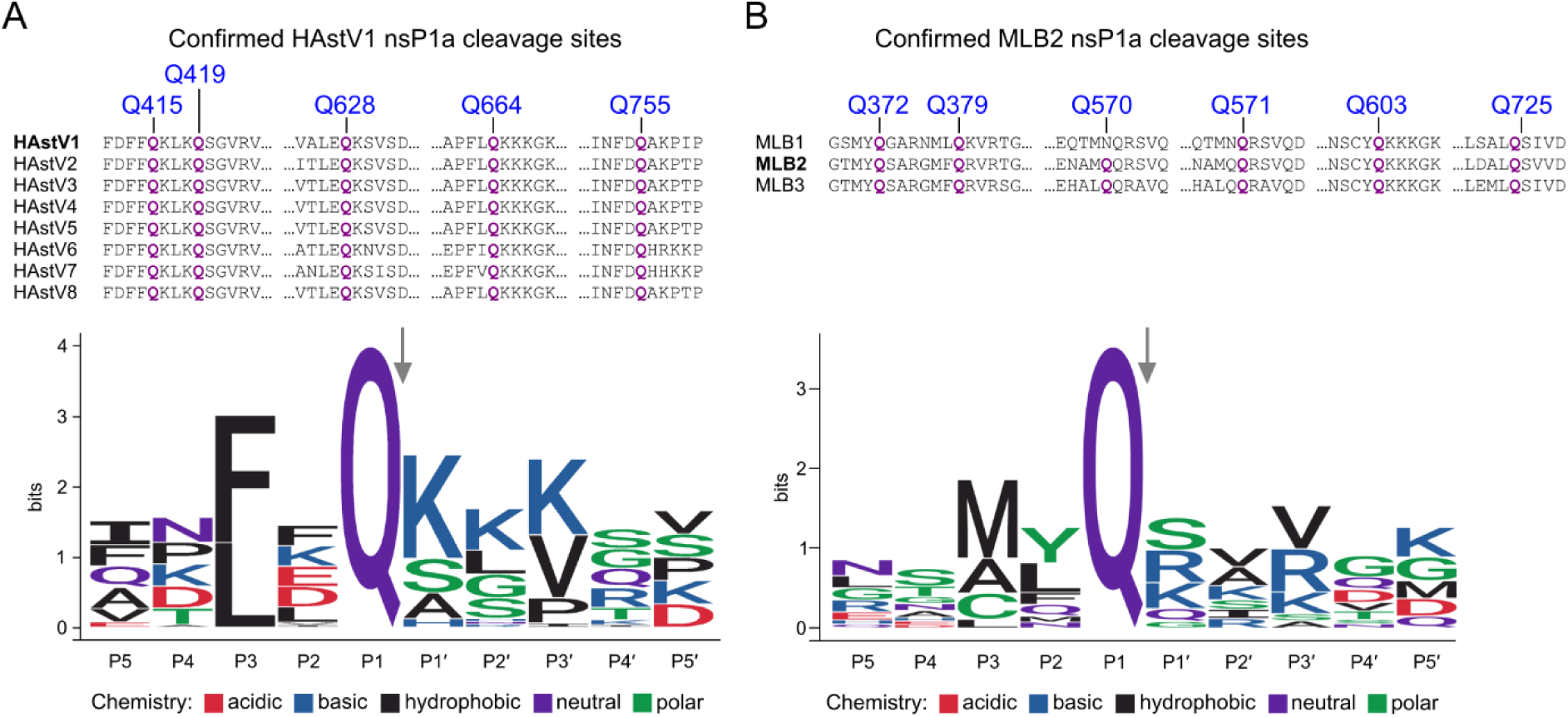
Cleavage site alignment and motif analysis between identified cleavage sites (P5 to P5′) for HAstV. (**A**) and MLB (**B**) sequences. Only the sequences from confirmed cleavage sites and closely related sequences were used for alignments and analyses.

## Discussion

Characterizing the processing of the astrovirus nonstructural polyprotein is crucial for understanding the viral life cycle, including the assembly of the replication complex and the dynamics of RNA replication, as well as for studying individual protein functions. We investigated both classical (HAstV1) and non-classical (MLB2) astrovirus nsP1a cleavage patterns using complementary experimental approaches: N-terminomics of virus-infected cells and detection of processing products from polyprotein overexpression in mammalian cells to find the boundaries of individual cleavage products.

Based on our experimental data, computational analyses, and previously reported findings^23^, we propose detailed cleavage maps of the astrovirus nonstructural polyproteins, identifying six final cleavage products for both HAstV1 and MLB2 nsP1a (Fig. 7A-B), which contrasts with the previously suggested four nonstructural proteins, nsP1a/1-nsP1a/4. We further update a consensus astrovirus genome map and the resulting proteins, mediated by a single viral and multiple host proteases – signal peptidases, trypsin, and caspases (Fig. 7C)^9,10,23,24,26–28^. Using our established replicon systems^9,24^, we demonstrated the possible functional relevance of these cleavage sites for viral replication. Mutations that significantly disrupt polyprotein processing also strongly impair replication activity in both HAstV1 and MLB2 replicon systems.

**Figure 7.**
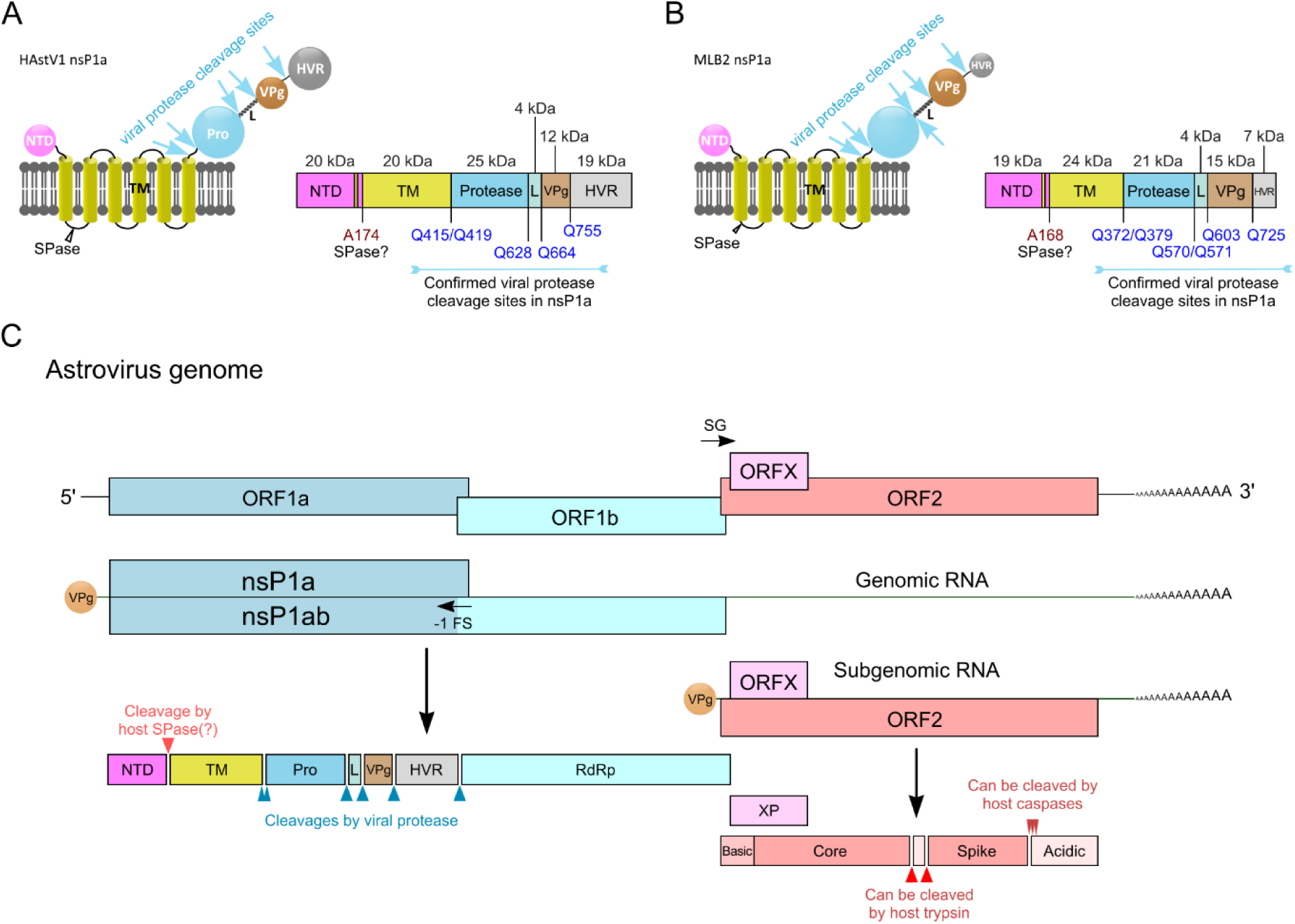
The map of HAstV1. (**A**) and MLB2 (**B**) astrovirus polyprotein processing. (**C**) Updated consensus map of the full astrovirus genome, based on identified HAstV1 and MLB2 cleavage sites and previously published data^9,10,23,24,26–28^. NTD, N-terminal domain; TM, transmembrane region; Pro, protease; L, linker region; VPg, viral protein genome-linked; HVR, hypervariable region; RdRp, RNA dependent RNA polymerase; ORF, open reading frame; SG, subgenomic promoter; –1 FS, (−1) frameshift site; SPase, signal peptidase.

Interestingly, some full cleavage blocks (Q628G and Q755G) were still tolerated in the HAstV1 replicon system. Whereas for MLB2, all fully blocked cleavages resulted in halted replication (Q372G+Q379G, Q571G, Q603G, and Q725G), highlighting differences in the requirements for polyporotein processing between classical and non-classical astroviruses.

The presence of two potentially redundant cleavage sites at the N-terminus of HAstV1 protease (Q415 and Q419) is intriguing. Overexpression of each mutant polyprotein individually does not affect the viral nsP1a processing patterns, suggesting that these residues do not control the order of processing. However, simultaneous mutation of both sites completely abolishes polyprotein processing (Fig. 3B). This effect is recapitulated in the replicon system, where the combined mutant exhibits a stronger effect on replicon activity (Fig. 5B). Similarly, processing of the MLB2 protease requires a dual N-terminal and dual C-terminal cleavages (Fig. 4B). These additional cleavages could be required for temporal and/or spatial control as different sites may become accessible or be cleaved at different stages of infection. Also, incomplete or alternative processing can lead to the production of various precursors that may have distinct functions from the fully mature or alternatively cleaved proteins^29–31^.

Polyprotein processing is tightly regulated in many (+)ssRNA viruses, ensuring the correct localization and function of replication complexes. This phenomenon is well characterized in picorna-, alpha-, flavi– and coronaviruses^32^. Polyprotein expression strategy enables the virus to condense its genetic material while allowing both temporal and spatial control of replicase activity, depending on the stage of polyprotein cleavage^33,34^. Understanding the structural reorganization of uncleaved and cleaved astrovirus polyproteins is critical for elucidating the mechanisms underlying replication complex assembly and regulation of viral RNA synthesis.

Interestingly, full protease processing is not required for the cleavage of HAstV1 VPg or HVR, as mutations at protease processing sites do not impair the release of VPg or HVR (Fig. 3B). This observation suggests that VPg release may occur early during infection, providing an essential RNA-binding protein that orchestrates both translation and replication. A similar mechanism has been observed in noroviruses, where multiple precursors containing the protease domain retain enzymatic activity and contribute to viral replication^29^.

While this study provides a comprehensive map of astrovirus polyprotein processing, several key questions remain to be addressed. Investigation of the sequential order of nsP1a cleavage events will offer valuable insights into the temporal regulation of viral replication^35,36^. The specificity of cleavages and functional differences between fully cleaved and precursor proteins remain to be determined. Characterisation of host signal peptidases (SPases) that cleave the NTD will complete the list of nsP1a cleavages. Taken together, the detailed characterization of the astrovirus polyprotein processing sites is important for understanding the organization of the viral genome and its replication strategy. Precise mapping of individual protein boundaries will enable in-depth functional and structural analyses, uncovering potential targets for developing antiviral therapeutics.

## Materials and methods

### Cells

HEK293T cells (ATCC) were cultured in DMEM supplemented with 1 mM L-glutamine, 10% fetal bovine serum (FBS), and antibiotics. Huh.7.5.1 (Apath, Brooklyn, NY) and Caco-2 (ATCC) cell lines were maintained in the same media supplemented with non-essential amino acids. BSR cell line was maintained in DMEM supplemented with 1 mM L-glutamine, 5% fetal bovine serum (FBS), and antibiotics. All cells were tested mycoplasma negative throughout the work (MycoAlert® Mycoplasma Detection Kit, Lonza).

### Plasmids and cloning

For mammalian expression of the HAstV1 and MLB2 nsP1a and its truncated variants, the coding sequence was inserted into vector pCAG-PM^37^ using *Afl*II and *Pac*I restriction sites. The resulting constructs were confirmed by sequencing. All mutations were introduced using site-directed mutagenesis and confirmed by sequencing.

For bacterial expression of folded domains of HAstV1 VPg (665-755 aa in ORF1a, L23513.1) and HVR (756-905 aa in ORF1b, L23513.1), the indicated coding sequence was PCR amplified and inserted into the T7 promoter-based pExp-MBP-TEV-CHis expression plasmid containing N-terminal MBP fusion tag, TEV protease cleavage site, and C-terminal 8×His-tag. The expression construct for the MLB2 VpG N-terminus (603-694) was prepared using the pExp-MBP-TEV-Mxe-CHis vector, where the protein-coding sequence was placed between MBP-TEV and 8×His-tagged Mxe GyrA intein.

Replicons were described earlier^9,24^. The resulting replicon plasmids were linearized with *Xho*I restriction enzyme prior to T7 transcription.

### Virus rescue, growth, and detection

HAstV1^9,38^ and MLB2^24^ were generated from infectious clones according to previously described protocols. The linearized infectious clones of HAstV1 (pAVIC1, L23513.1) and MLB2 (pMLB2, ON398706) were used as templates to produce capped T7 RNA transcripts using T7 mMESSAGE mMACHINE Transcription kit (Invitrogen) according to the manufacturer’s instructions. For virus recovery, Huh7.5.1 cells were used for MLB2 and BSR cells for HAstV1. Cells were electroporated with 20 µg T7 RNA in 800 µl PBS with two pulses at 800 V and 25 µF using a Bio-Rad Gene Pulser Xcell electroporation system. The passaging of MLB2 was performed in Huh7.5.1 cells and passaging of HAstV1 in Caco-2 cells. For HAstV1, trypsin at a final concentration of 0.7 µg/ml was added during the infection. Both viruses were titrated by immunofluorescence using capsid-specific antibodies^9,24^.

### Proteomics

Caco-2 cells were infected at MOI 1 with HAstV1 and protein lysates collected at 20 hpi in lysis buffer consisting of 1% SDS, 2× Thermo HALT protease inhibitor, 100 mM HEPES pH 8 and 1% NP40. Huh7.5.1 cells were infected with MLB2 at MOI 1 and lysed at 40 hpi. Samples were processed as per the published TMT-HUNTER protocol^39^ with modifications. Briefly, cell lysates were heated to 95 °C for 5 min, then cooled to room temperature and treated with Benzonase Nuclease (Sigma Aldrich) at 37 °C for 30 min. Protein concentrations were normalized using lysis buffer. Dithiothreitol (DTT) was added to 4 mM and samples heated to 60 °C for 10 min. Iodoacetamide (IAM) was added to 14 mM for cysteine alkylation at room temperature for 30 min in the dark, then IAM was quenched by adding 3 mM DTT and incubating at room temperature for 20 minutes. Samples were cleaned up using SP3 method^40^. Next, the beads with the samples bound were resuspended in 40 µl of 4 M guanidine hydrochloride containing 10 mM Tris(2-carboxyethyl)phosphine (TCEP) in 250 mM HEPES pH 8 was added to each sample for solubilization of precipitated proteins. Samples were sonicated in water bath for 5 minutes and agitated at 1,300 rpm, room temperature, for 30 min to disaggregate beads. TMTpro tags – 0.5 mg per sample – were solubilized by sequential addition of 10 µl anhydrous acetonitrile (MeCN) and 50 µl of anhydrous dimethyl sulfoxide (DMSO), then transferred into the samples microtubes and incubated at room temperature, 1,300 rpm, for 1.5 h. TMTpro channels allocation to samples was randomized using the function sample() in R. Excess of TMTpro was quenched by adding hydroxylamine to 0.4% w/v and incubating for 45 min at room temperature. Samples were pooled together, and SP3-cleaned. Precipitated beads-protein aggregates were resuspended in 200 mM HEPES pH 8 to 1 mg/ml final protein concentration and water-bath sonicated for 5 minutes. The resuspended sample was divided equally for parallel digestion with Trypsin Gold (Promega) and Chymotrypsin Endoproteinase (Thermo Scientific) at 1:50 enzyme-to-protein ratio. Chymotrypsin digestion was supplemented with 1 mM calcium chloride, and both digestion tubes were incubated at 37 °C, 1,300 rpm, for 16 h. Digests were centrifuged at 15,000× g for 10 minutes, placed in a magnetic rack, and the supernatants were transferred into clean tubes. To each digest, undecanal was added to 1:40 peptide to undecanal ratio, plus sodium cyanoborohydride to 30 mM and ethanol to 40 % v/v final concentration. The samples were water-bath sonicated for 2 min, and incubated at 37 °C for 1 h, 1,300 rpm. Then, undecanal-treated trypsin and chymotrypsin digests were acidified to 0.5% v/v TFA and loaded separately to C18-Macrospin columns (Nest group) previously equilibrated in 0.1% TFA in 40 % ethanol. The flowthroughs enriched for N-terminal peptides were collected, dried in a vacuum centrifuged, desalted on C18-Macrospin columns and dried down again before off-line basic reverse phase (bRP) fractionation. Off-line bRP and LC-MS/MS analysis of concatenated bRP fractions were performed for trypsin and chymotrypsin samples as described previously^39^.

Spectral data was searched against Uniprot Human reviewed sequences (download: 30/05/2023) plus the six-frame translation of HAstV1 and MLB2 ORFs, using Fragpipe v20.0 with MSFragger v3.8^41^ and Philosopher v5.0^42^. The TMTpro workflow was selected and modified for N-terminomics analysis. In brief, for samples digested with trypsin, specificity was set to “trypsin_r (P1 = R, P1’ ≠ P)” and up to 1 missed cleavage site was allowed, and for chymotrypsin samples, the enzyme specificity was “P1 = FLWYM, P1’≠ P) and up to 3 missed cleavage sites allowed. In both cases, digestion mode was set to “semi-specific”. Variable modifications included protein N-terminal acetylation, methionine oxidation, and peptide N-terminal TMTpro or Gln/Glu to pyroglutamate as variable modifications. Cysteine carbamidomethylation and lysine TMTpro were set as fixed modifications. Data was filtered at 1% FDR at the PSM, peptide, and protein levels. Peptide identifications from both trypsin-and chymotrypsin-digested samples were filtered to retain viral peptides N-terminally blocked by TMTpro, and peptides with trypsin– or chymotrypsin-like N-terminal cleavages were removed. The mass spectrometry data has been submitted to proteomeXchange via the PRIDE repository and has accession number PXD058484.

### Overexpression of astrovirus nsP1a

HEK293T cells were transfected with 1:2 ratio of pCAG-nsP1a overexpression constructs (µg) to Lipofectamine 2000 (µl). Protein cell lysates were made using 1× radioimmunoprecipitation assay (RIPA) buffer and 1× Halt protease inhibitor cocktail. Cells were lysed 24 hours post transfection and analyzed by western blotting.

### SDS-PAGE and immunoblotting of virus-infected cells

Cells were infected with MLB2 and HAstV1 (MOI 1). Cells were lysed at the indicated time points, and proteins were separated on a 10-15% SDS-PAGE before being transferred to 0.2 µm nitrocellulose membrane using Trans-Blot Turbo Transfer System semi-dry transfer. Primary antibodies at the following concentrations were used: HAstV1 protease (custom, 1:2000), MLB2 protease (custom, 1:500), HAstV1 VPg (custom, 1:1000), MLB2 VPg (custom, 1:1000), HAstV1 HVR (custom, 1:1000) and anti-HA tag (Roche, 11867423001, 1:1000)), and anti-tubulin (Abcam, ab6160 and ab15568, 1:3000). Secondary antibodies (Licor IRDye 800 and 680) were added at 1:3000 dilution for IR-based detection. Immunoblots were imaged on a LI-COR ODYSSEY CLx imager and analyzed using Image Studio version 5.2.

### Purification of His-tagged astrovirus proteins and generation of specific antibodies

The HAstV1 VPg and HVR proteins were produced in Rosetta 2 (DE3) cells (Novagen) cultured in 2×YT media with overnight expression at 18 °C induced with 0.4 mM IPTG. The proteins were purified first by immobilized metal affinity chromatography using PureCube Ni-NTA resin and then by affinity chromatography using amylose resin (NEB). MBP fusion tag was removed by the cleavage with TEV protease (produced in-house). Proteins were further purified by heparin chromatography using HiTrap Heparin HP or anion-exchange chromatography using HiTrap QHP 5 ml column (Cytiva) and, finally, by size exclusion chromatography using a Superdex 200 16/600 column (Cytiva). Protein solution in 50 mM Na-phosphate pH 7.4, 300 mM NaCl, 5% glycerol was concentrated to 2 mg/ml and used for immunization.

The MLB2 VPg (N-terminal part) was produced in Evo21(DE3) cells and purified first by Ni-affinity chromatography, and the C-terminal intein was activated by the addition of 50 mM β-mercaptoethanol. Then the N-terminal MBP fusion was removed by TEV protease cleavage, and the proteins were additionally purified by cation-exchange on HiTrap SP HP 5 ml column (Cytiva) and size-exclusion chromatography runs. The resulting protein in PBS with 10% glycerol was concentrated to 1 mg/ml and used for immunization.

Antibodies against indicated proteins were generated in rabbits using 5-dose 88-day immunization protocol. Sera were used for specific affinity purification, followed by purification of specific IgG fractions (BioServUK Ltd).

### Replicon assays

Linearized replicon-encoding plasmids were used as templates to generate T7 RNAs using T7 mMESSAGE mMACHINE Transcription kit according to manufacturer’s instructions. The replicon RNA was purified using Zymo RNA Clean & Concentrator kit and quantified. Cells were transfected in triplicate using the previously described reverse transfection protocol^24^. Three independent experiments, each in triplicate, were performed to confirm the reproducibility of the results.

### Protein structure prediction

The 4 kDa (L) protein 3D structure was predicted using AlphaFold 3^43–45^. The confidence of prediction was quantified by pLDDT, the predicted local distance difference test on the Cα atoms, with >90% for most regions.

### Sequence alignments

Cleavage sites for HAstV and MLB sequences were aligned using ClustalW^46^. The following sequences were used for analysis: HAstV1 (L23513.1), HAstV2 (MK059950.1), HAstV3 (MK059951.1), HAstV4 (MK059952.1), HAstV5 (MK059953.1), HAstV6 (GQ495608.1), HAstV7 (MK059955.1), HAstV8 (AF260508.1), MLB1 (ON398705.1, FJ222451.1), MLB2 (ON398706.1, JF742759.1), and MLB3 (NC_019028.1, MT766337.1).

### Statistical analyses

Data were graphed and analyzed using GraphPad Prism. Where appropriate, data were analyzed using one-way ANOVA test. Significance values are shown as \*\*\*\**p* < 0.0001, \*\*\**p* < 0.001, \*\**p* < 0.01, ns – nonsignificant.

### Data availability

The mass spectrometry proteomics data have been deposited to the ProteomeXchange Consortium (http://proteomecentral.proteomexchange.org) via the PRIDE partner repository^47^ with the dataset identifier PXD058484.

## Acknowledgments

This work was funded by a Sir Henry Dale Fellowship (220620/Z/20/Z) from the Wellcome Trust and the Royal Society to VL. EE is funded by an Academy of Medical Sciences Springboard Award (SBF006\1008) supported by the British Heart Foundation, Diabetes UK, the Global Challenges Research Fund, the Government Department for Business, Energy and Industrial Strategy and the Wellcome Trust, the Medical Research Council [MR/X000885/1], BBSRC [BB/W019744/1] and a Wellcome Trust Career Development Award [227831/Z/23/Z]. DN is supported by the Elizabeth Mann-funded studentship from the Department of Pathology. For the purpose of Open Access, the authors have applied a CC BY public copyright license to any Author Accepted Manuscript (AAM) version arising from this submission.

The authors declare that they have no conflict of interest.

